# IN VIVO ANALYSIS OF THE ROLE OF GASDERMIN-B (GSDMB) IN CANCER USING NOVEL KNOCK-IN MOUSE MODELS

**DOI:** 10.1101/2021.05.27.445936

**Authors:** David Sarrio, Alejandro Rojo-Sebastián, Ana Teijo, María Pérez-López, Eva Díaz-Martín, Lidia Martínez, Saleta Morales, Pablo García-Sanz, José Palacios, Gema Moreno-Bueno

## Abstract

**Background:** Gasdermin-B gene (GSDMB) is frequently over-expressed in tumors, and its shortest translated variant (isoform 2; GSDMB2) increases aggressive behavior in breast cancer cells. Paradoxically, GSDMB could have either pro-tumor or tumor suppressor properties depending on the biological context. Since GSDMB gene is not present in the mouse genome, deciphering fully the functional roles of GSDMB in cancer requires novel *in vivo* models.

**Methods:** We first generated by gene targeting a conditional knock-in mouse model (R26-STOP-GB2) harboring human GSDMB2 transcript within the *ROSA26* locus. We next derived the R26-GB2 model ubiquitously expressing GSDMB2 in multiple tissues (confirmed by western blot and immunohistochemistry) and performed a comprehensive histopathological analysis in multiple tissues from 75 male and female mice up to 18 months of age. Additionally, we produced the double transgenic model R26-GB2/MMTV-PyMT, co-expressing GSDMB2 and the Polyoma-Middle-T oncogene, and assessed breast cancer generation and progression in GSDMB2-homozygous (n=10) and control (n=17) female mice up to 15 weeks of age.

**Results:** In the R26-GB2 model, which showed different GSDMB2 cytoplasmic and/or nuclear localization among tissues, we investigated if GSDMB2 expression had intrinsic tumorigenic activity. 41% of mice developed spontaneous lung tumors, but neither the frequency nor the histology of these neoplasias was significantly different from wildtype animals. Strikingly, while 17% control mice developed gastric carcinomas, no GSDMB2-positive mice did. No other tumor types or additional histological alterations were frequently seen in these mice. In the R26-GB2/MMTV-PyMT model, the strong nucleus-cytoplasmic GSDMB2 expression in breast cancer cells did not significantly affect cancer formation (number of tumors, latency, tumor weight, histology or proliferation) or lung metastasis potential compared to controls.

**Conclusions:** GSDMB2 expression alone does not have an overall tumorigenic potential in mice, but it might reduce gastric carcinogenesis. Contrary to human cancers, GSDMB2 upregulation does not significantly affect breast cancer generation and progression in mouse models. However, to evidence the GSDMB functions in cancer and other pathologies *in vivo* may require the presence of specific stimulus or cellular contexts. Our novel mouse strains will serve as the basis for the future development of more precise tissue-specific and context-dependent cancer models.

## BACKGROUND

The Gasdermins (GSDMs, named after their Gastric and Dermal expression) are cytosolic proteins of around 50 KDa [1, 2] that have been functionally involved in the genesis and development of cancer and multiple diseases (reviewed in [3, 9]). The GSDM family comprises six genes in the human genome [1, 2]: *GSDMA* and *GSDMB* (which are both located in the 17q21.1 region), *GSDMC* and *GSCMD* (in 8q24); *GSDME/DFNA5* (7p15.3) and *DFNB59/PJVK* (2q31.2). Mice have ten GSDM genes but *GSDMB* gene (also known as *GSDML* and *PRO2521*) is the only GSDM member that is not present in the mouse or rat genomes [1, 2].

The diverse biological functions of GSDM proteins have started to emerge recently, and multiple studies indicate that each family member, except possibly DFNB59, can produce cell death, through specific mechanisms including pyroptosis (lytic and pro-inflammatory cell death), apoptosis, mitochondrial damage or autophagy (reviewed in [3-5, 10, 11]). These cell-death promoting functions are normally auto-inhibited through the intramolecular interaction of GSDMs N-terminal (NT) and C-terminal (CT, inhibitory) domains. Under certain stimuli and circumstances, the NT is exposed or released, mostly via specific protease cleavage and produces cell damage generally through the formation of NT membrane pores [12-16], among other potential mechanisms [10, 17, 18]. In the recent years, these GSDM pro-cell death activities have been proposed to play a role in the pathogenesis of multiple diseases (inflammatory, infectious, neurological, among others [3-5]) and also in the progression and clinical behavior of cancer [6-9, 19]. In this sense, due to their potential cytotoxic function, GSDME and GSDMA are generally silenced in cancers and broadly considered as potential tumor suppressor genes [7, 9, 19-21], while GSDMC and GSDMD have been associated to pro- and anti-tumor effects, depending on the context [22-26]. Strikingly, GSDMB is frequently over-expressed in the cytoplasm (and/or the nucleus) of esophageal, gastric, colon, liver and breast tumor cells, as well as cervical and head and neck squamous cell carcinomas [27-33]. Moreover, GSDMB upregulation associates with poor prognosis and/or aggressive behavior in breast and other cancer types [28, 30, 32-34], suggesting a role as a potential oncogene [27-33]. In fact, our previous work have demonstrated that GSDMB overexpression increases invasive and metastatic behavior in breast cancer cells without affecting cell proliferation [32-34]. Moreover, GSDMB overexpression, mostly due to gene amplification, occurs in more than 60% of HER2+ breast carcinomas [33] and in at least 25% of HER2+ gastric cancers [27]. In HER2+ breast cancer high GSDMB levels associate with metastasis, poor prognosis and resistance to anti-HER2 therapies [33, 34]. Thus, GSDMB upregulation promotes multiple pro-tumor functions, and can be a novel therapeutic target in cancer. In fact, our lab has demonstrated that multiple pro-cancer functions can be reduced *in vitro* and *in vivo* by the intracellular delivery of a GSDMB antibody through biocompatible nanocapsules, into HER2+/GSDMB+ breast cancer cells [34]. Paradoxically, GSDMB intrinsic cytotoxic function could also be activated in tumor cells, via lymphocyte-derived Granzyme A (GZMA) cleavage of GSDMB-NT [35] or by an anti-GSDMB therapeutic antibody [34].

Additionally, the existence of at least four GSDMB protein isoforms, adds more complexity into the biological roles of GSDMB. The alternative usage of exons 6 and 7 (encoding residues located in the protein inter-domain [36]) can lead to the expression of either GSDMB isoform 3 (NM_001165958.1; “ full-length” protein of 416 aminoacids –aas-and 47.4 KDa); isoform 1 (NM_001042471.1; lacks exon 6, 403 aas and 45.8 KDa); isoform 2 (NM_018530.2; lacking both exons 6 and 7, 394 aas and 45 KDa) or isoform 4 (NM_001042471.1; does not have exon 7, 407 aas and 46.5 KDa). The differential expression of these isoforms in cells and tissues could lead to distinct functional consequences in normal and pathological contexts, such as inflammatory diseases [37-39] and cancer [29, 30, 32, 40]. In this sense, we previously showed that overexpression of either isoform 1 or 2 increases motility and invasion *in vitro*, but only the isoform 2 (the shortest transcript; hereafter referred to as GSDMB2) promotes tumor growth and metastasis when MCF7 breast cancer cells were xenografted in immunocompetent mice [32]. This data suggested an increased pro-tumor potential for GSDMB2 isoform.

To elucidate the roles of GSDMB in cancer development and progression *in vivo*, here we first generated by gene targeting technology a conditional mouse model (R26-STOP-GB2) harboring human GSDMB2 transcript and GFP within the *ROSA26* locus. Then, we activated the expression of GSDMB2 in all mice tissues, mimicking human *GSDMB* expression, which occurs in multiple organs and tissues, including the digestive tract and liver [27, 29, 30], lymphocytes [41], lung epithelia [37, 41], among others [5, 6]. In this mouse model, termed R26-GB2, we performed the first comprehensive *in vivo* study of GSDMB effects on tumor initiation and development.

## METHODS

### Animal models

The commercial mouse strains B6.FVB-Tg (EIIa-Cre)C5379Lmgd/J and FVB/N-Tg(MMTV-PyVT)634Mul/J were purchased from JaxMice. The FVB/NCrl strain was purchased from Charles River. The generation of three novel animal models (R26-STOP-GB2, R26-GB2, and R26-GB2/MMTV-PyMT) is described below. All mouse studies were performed in agreement with the procedures and protocols approved by the committees of ethical and animal welfare as detailed in the Ethics approval section.

### Generation of conditional GSDMB2 Knock-in mice (R26-STOP-GB2)

We generated, in collaboration with the CNIO Transgenic mice Service, mice harboring human GSDMB isoform 2 transcript (NM_018530.2) essentially as previously reported [42, 43]. Using gene-targeting technology, we inserted by homologous recombination into the endogenous *ROSA26* (R26) locus [43] a construct containing a loxP-flanked PGK-neomycin-STOP cassette followed by the human GSDMB isoform 2 cDNA (GSDMB2) fused with the HA-tag sequence [32] (**Fig. 1A**). The construct also contains an IRES-sequence followed by the Green Fluorescent Protein (GFP) gene reporter, which helps in the identification of the Knock-in (KI) animals (**Fig. 1A**). Expression of the construct is under the control of endogenous *Rosa26* promoter, which allows ubiquitous and moderate levels of expression of the transgene [43]. The targeting vector for the homologous recombination was generated by the Gateway cloning DNA technology using the pEntry plasmid harboring GSDMB2-HA cDNA and the pROSA26-DV vector [43], as reported previously [42]. Recombinant clones were sequence-verified using the S1F (5’-ATCATGTCTGGATCCCCATC-3’) and S2R (5’-GGGGCGGAATTCGATATCAAG-3’) primers. The targeting construct was then electroporated into ES cells (C57BL6×129 background; CNIO Transgenic Mice Service), and positive ES clones harboring the construct in the correct orientation were detected by diagnostic PCR (conditions detailed in **Supplementary data: Table S1**). Two positive ES clones were used for aggregation with CD1 embryos (CNIO Transgenic mice Service) obtaining 18 male chimeras (all >80% chimerism). After crossing with FVB wildtype (WT) female mice, the correct transmission of the transgene was demonstrated by two PCR tests (**Fig. 1 B, C**): a) insertion of the transgene into 3’-and 5’-arm; b) presence of GSDMB-HA transgene and GFP gene. PCR conditions are detailed in **Supplementary data: Table S1**; Uncropped PCR gels are provided in **Supplementary data: Fig. S1**. After confirmation of correct transgene insertion by PCR, we derived one mouse strain of conditional GSDMB2 expression, referred to as R26-STOP-GB2. This strain has mixed genetic background (C57BL6×129 from the ES, CD1 from the embryo aggregation and crossed twice with FVB).

**Fig. 1.**
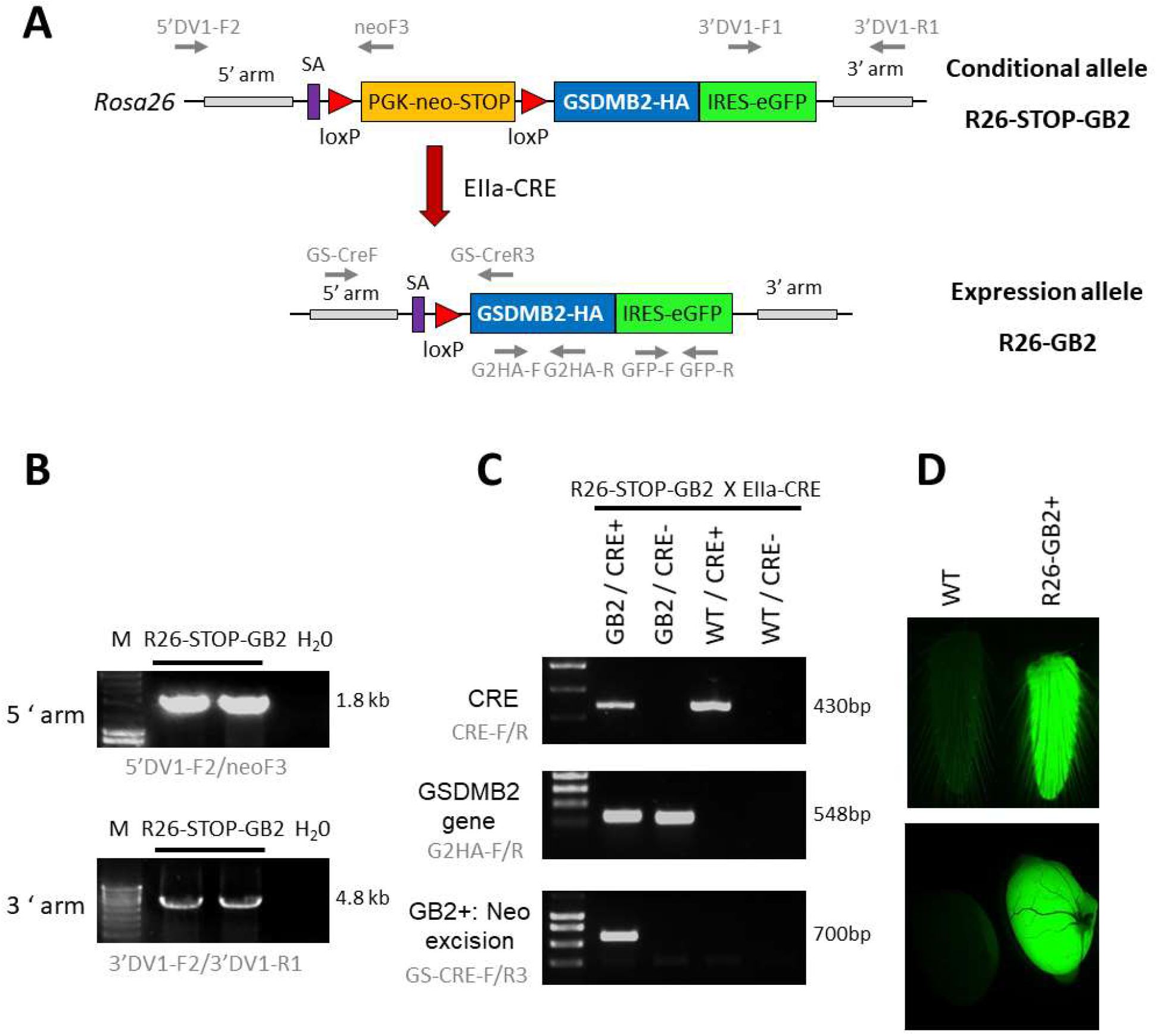
Generation of knock-in mouse models harboring human GSDMB isoform 2 transcript within the ROSA26 locus. **A:** Schematic representation of the GSDMB isoform 2 (GB2) targeted floxed allele (top, conditional model) and the expression allele (bottom) within the ROSA26 (R26) locus. The construct contains a splicing acceptor signal (SA), the PGK-neomycin-STOP cassette flanked by LoxP sites, the human GSDMB2 isoform 2 cDNA sequence (GB2) fused with HA tag, followed by the IRES-GFP reporter gene. After crossing with EIIa-Cre strain (red arrow), the Cre-mediated excision of the PGK-neomycin-stop element allows the ubiquitous expression of the GB2-HA/GFP tandem under the control of the ROSA26 promoter (R26-GB2). The primer pairs for PCR analyses are also detailed (gray arrows). **B:** Diagnostic PCR analysis of positive ES cell clones showing the proper insertion of the recombinant R26-STOP-GB2 allele. H20, Negative control. **C:** Examples of genotyping PCR analysis (primers in gray), demonstrating the excision of neo-stop cassete in Cre+ /GB2 mice. **D:** Ubiquitous expression of the transgenes is verified by GFP fluorescent emission of fresh tail skin (top) and testes (bottom) from WT and GSDMB-positive R26-GB2 mice. Full-length gels are presented in Supplementary data: Fig. S1.

### Generation of Knock-in mice with ubiquitous expression of GSDMB2-HA (R26-GB2)

To analyze *in vivo* the consequences of ubiquitous GSDMB2 expression in mice, conditional R26-STOP-GB2 male animals were crossed with female mice of the B6.FVB-Tg (EIIa-Cre)C5379Lmgd/J strain (JAXmice). Adenoviral EIIa promoter expression is restricted to oocytes and preimplantation stages of the embryo, and thus Cre-mediated recombination occurs in a wide range of tissues, including the germ cells that subsequently transmit the genetic modification to progeny. The deletion of the neo-STOP cassette by Cre permits the transcriptional expression, mediated by R26 promoter, of the bicistronic mRNA GSDMB2-HA-IRES-GFP (**Fig. 1A**). To verify the correct excision of the neo cassette and the subsequent activation of the transgenes we designed a diagnostic PCR reaction that preferentially amplifies the excised allele in DNA obtained from tail skin (**Fig. 1C**). PCR conditions are detailed in **Supplementary data: Table S1**. Uncropped PCR gels are provided in **Supplementary data: Fig. S1**. Moreover, GB2-HA-GFP expression was demonstrated by GFP fluorescence imaging in the tail and other organs (**Fig. 1D**) using a Leica MZ10F magnifier. After validation of the transgenes ubiquitous expression we crossed heterozygous animals to remove the Cre recombinase and to obtain a line expressing germline GSDMB2-HA-GFP in all tissues. This mouse model, named R26-GB2, with mixed background was crossed two times with the FVB/NCrl strain (Charles River) to ensure that it contained at least 50% FVB genetic background.

### Generation of a breast cancer mouse model expressing PyMT oncogene and GSDMB2-HA (R26-GB2/MMTV-PyMT)

In order to study the effect of GSDMB2-HA in breast cancer progression, the R26-GB2 model was crossed with the FVB-MMTV-PyMT strain (JaxMice). The mammary glands of female animals from the MMTV-PyMT express the Polyoma Middle T antigen under the regulation of the MMTV (Mouse Mammary Tumor Virus) promoter [44]. These mice develop spontaneous invasive breast carcinomas and metastatic lung colonization by 14 weeks of age [44]. To generate R26-GB2/MMTV-PyMT double transgenic animals, male homozygous MMTV-PyMT mice were crossed with female GSDMB2 homozygous R26-GB2. Then, male mice heterozygous for PyMT and GSDMB2 were crossed with female GSDMB2 heterozygous R26-GB2 animals. From the resulting offspring, we selected female mice heterozygous for PyMT and either GSDMB2 homozygous or WT for the breast cancer tumorigenesis assays. Genotyping of the PyMT oncogene was performed as described in **Supplementary data: Table S1**. The R26-GB2/MMTV-PyMT model derives from three crossings with mice of the FVB genetic background.

### Phenotypic and histological characterization of R26-GB2 model: study of spontaneous tumorigenesis

Heterozygous R26-GB2 animals were crossed to obtain at least 18 animals from each of the genotypes. A total of 80 mice (42 males, 38 females) corresponding to the three GB2 genotypes (WT, n=27; GB2+/-, n=34 and GB2+/+, n= 19) were studied up to 18 months of age. Five animals died spontaneously and no necropsy could be performed, thus were excluded from the histological analyses. Mice were monitored weekly for the appearance of tumor masses (in any part of the body) or other pathological signs (outcome). Animals were sacrificed when they reached 18 months of age or showed any of the criteria for early termination (scored as 4) specified in Orellana-Muriana (2013) [45]. These criteria include, tumors >15mm, ulceration or infection of the tumors, body weight loss >20%, enlarged lymph nodes, or extensive skin ulceration, among others. Animals were euthanized in a CO_2_ chamber (fill rate of 30% of the chamber volume per minute) and necropsy was performed immediately. We extracted all organs with macroscopic signs of cancer or other pathologies, as well as other selected organs with normal appearance. Tissues were fixed in 10% formalin and embedded in paraffin blocks. For histologic examination, Hematoxylin-Eosin stained tissue sections were analyzed by two Pathologists (ARS and AT). A total of 328 tissue sections were reviewed (median= 3 organs per mice; minimum 1 and maximum of 14).

### Study of breast tumorigenesis and metastatic potential in R26-GB2/MMTV-PyMT mice

Twenty-seven female mice of the R26-GB/MMTV-PyMT model were used. All animals were heterozygous for PyMT and either homozygous for GSDMB2 (n=10) or WT (n=17). The appearance of breast tumors was monitored 3 times a week until animals reached 15 weeks of age (endpoint). Then, we analyzed the cancer incidence, latency (time until detection of palpable tumor), number and weight of tumors. For this, mice were sacrificed by CO_2_ inhalation (fill rate of 30% of the chamber volume per minute) and all mammary glands with breast tumors were extracted (mean 5.6 tumors/mice), washed in 1x PBS, measured with caliper and weighted on a scale. The biggest tumor of each mice was selected and half of the cancer tissue was processed for subsequent histological analysis by two pathologist (as described above). The rest of the tissue was quickly frozen in dry-ice and stored at -80°C. Moreover, to assess metastatic potential the whole lungs were extracted and processed for subsequent paraffin-embedding. Then, whole lungs were serially sectioned into 5µm-thick tissue sections using a microtome (Leica RM 2255). From these slides, we selected four sections separated by 100 µm in depth. Thus, the analyses of these four combined sections covered > 400 µm in depth. The sum of metastatic foci observed in the four slides was calculated. Metastatic lesions that appear repeatedly in two or more slides were counted once.

### Immunohistochemical analysis

GSDMB2-HA expression was analyzed by immunohistochemistry in 5µm-thick tissue sections using rat anti-HA (1:200; 3F10, ROCHE) or mouse monoclonal anti-GSDMB (1:10, [33]), following standard methods. Tumor proliferation in the R26-GB2/MMTV-PyMT animals was assessed by PCNA (proliferating cell nuclear antigen) immunostaining using the MAB424R antibody (1:10,000; clone p10, Millipore). Briefly, after an antigen-retrieval step (Leica Bond ER solution-1, citrate buffer 10mM pH 5.9-6.1) the primary antibodies were incubated for 1h at RT, followed by secondary-HRP antibody incubation. The staining was revealed by DAB standard Leica procedure. In negative controls, the primary antibodies were omitted. Immunohistochemical images were taken from representative samples with an Axiophot (Zeiss) microscope coupled with a color DP70 (Olympus camera), using the Olympus DP controller software. For immunofluorescence analysis, secondary goat anti-rat IgG-Alexa 547 (1:1000, Molecular probes) was incubated for 1h at RT. Slides were stained with 1:10,000 DAPI (4’,6-diamino-2-fenilindol, Molecular Probes), mounted with Prolong Diamond Antifade Mountant (Molecular Probes) and analyzed by confocal microscopy (LSM710, Zeiss).

### Analysis of GSDMB2-HA-GFP expression in tissues by western blot (WB)

Six R26-GB2 mice (3 males, 3 females) from each of the GB2 genotypes (WT, GB2+/-, GB2+/+) were sacrificed at 20 weeks of age. Sixteen organs were removed, chopped and immediately stored at -20°C. Tissues were homogenized in 50-200 µl lysis buffer (0.1M NaCl, 0.05M Tris HCl pH 7.9, 5μM MgCl_2_, 5μM CaCl_2_, 2% SDS supplemented with 1x protease inhibitor cocktail, ROCHE) by sonication on ice. Lysates were clarified by centrifugation (10.000 rpm, 5 min) and quantified by the BCA method (Pierce). Fifty µg of total proteins/per sample were loaded on 10% SDS-PAGE gels. Western blots were performed by standard methods using rat anti-HA (1:1000; clone 3F10, ROCHE), rabbit anti-GFP (1:2000; A6455, Molecular Probes) and mouse anti-GAPDH (1:50,000; 6C5, Calbiochem). As positive control, all rounds of WB contained a sample of MCF7 cells expressing GSDMB2-HA [32] and GFP constructs.

### Flow Cytometry

To evaluate the proportion of white blood cells from R26-GB2 mice expressing the GSDMB-HA-GFP transgenes we analyzed GFP emission by Flow Cytometry (Cytomics FC 500MPL, Beckman Coulter). Total leukocyte cells, not any specific subpopulation, were analyzed. Peripheral blood from WT and GB2+/+ mice was extracted and processed following the method reported before [46].

### Statistical analyses

Data was obtained from all available animals in the study (each mouse corresponds to a data point) and, unless otherwise specified, no data points were excluded from the analyses. The normal distribution of the continuous data was confirmed by the Kolmogorov–Smirnov test. Statistical analyses were performed using GraphPad 6.0 (GraphPad Software, Inc.) using Chi^2^ or Fisher’s exact tests to assess differences in categorical variables, and ANOVA or Student t-test for continuous variables. A *p value* <0.05 was considered as statistically significant.

## RESULTS

### Generation of the knock-in mouse model ubiquitously expressing GSDMB2-HA (R26-GB2)

To assess *in vivo* the functional role of GSDMB in tumorigenesis and cancer progression we selected the gene isoform 2 (GSDMB2) since the over-expression of this transcript promotes invasive and metastatic behavior of MCF7 breast cancer cells [32]. As GSDMB is the only GSDM gene not present in the mouse genome [1,2], using gene-targeting technology, we generated a KI model (named R26-STOP-GB2) harboring, within the endogenous *ROSA26* locus, the human GSDMB2 cDNA fused with the HA-tag and followed with the GFP transgene (**Fig. 1A,B**). After crossing with the EIIa-Cre strain, the ubiquitous expression of Cre-recombinase produced the excision of the Neomycin-STOP cassette (**Fig. 1A, C**), allowing the transcriptional activation, by the endogenous R26 promoter, of GSDMB-HA and GFP transgenes in the whole body of the animal. In these mice, GFP light emission, used as a readout of transgene expression, was clearly detected in some fresh tissues, such as tail skin or testes (**Fig. 1D**). Since EIIa-Cre-mediated recombination occurs also in the germ cells, that subsequently transmit the genetic modification to progeny, we crossed heterozygous animals to remove the Cre recombinase and to obtain a second line, named R26-GB2, expressing germline GSDMB2-HA and GFP transgenes in all tissues.

Homozygous (hereafter referred to as GB2+/+) and heterozygous (GB2+/-) mice from this model are viable and fertile, they reproduce normally, and female mice can nurse their litter normally. In crossings between heterozygous animals, the transgene is transmitted with expected frequencies of the Mendelian inheritance. Transgenic mice do not show evident morphological and developmental alterations or signs of abnormal behavior. GB2+/+ mice tend to have slightly higher body weight, especially in males, than WT animals but the differences do not reach statistical significance (**Supplementary data: Fig. S2**).

### Expression and intracellular localization of GSDMB2-HA in tissues

First, by Western Blot we verified in male and female GB2 mice the specific expression of GSDMB2-HA and GFP proteins in 14 different organs (**Fig. 2A**). Expression of the transgenic construct in peripheral blood leukocytes was additionally demonstrated by WB and FACs analysis (**Fig. 2B,C**), where more than 90% of cells showed GFP expression (**Fig. 2C**). GSDMB is a cytoplasmic protein although nuclear expression has been also reported in certain normal tissues, tumors and cancer cell lines [29, 30, 32]. Thus, we next analyzed GSDMB2-HA expression and subcellular localization in diverse GB2+/+ and WT tissues by immunohistochemistry using an anti-HA antibody. GSDMB2 showed mainly cytoplasmic localization in some tissues, such as breast, pancreas or liver, while nucleo-cytoplasmic staining was typically seen in specific tissues/cell types (**Fig. 3, Supplementary data: Figure S3 and Table S2**). Nuclear staining was particularly strong in the squamous epithelia of the esophagus, skin epidermis, hair follicles and sebaceous glands, as well as colon epithelia (**Fig. 3)** and testicles (**Supplementary data: Figure S4**), among others (**Supplementary data: Table S2**).

**Fig. 2.**
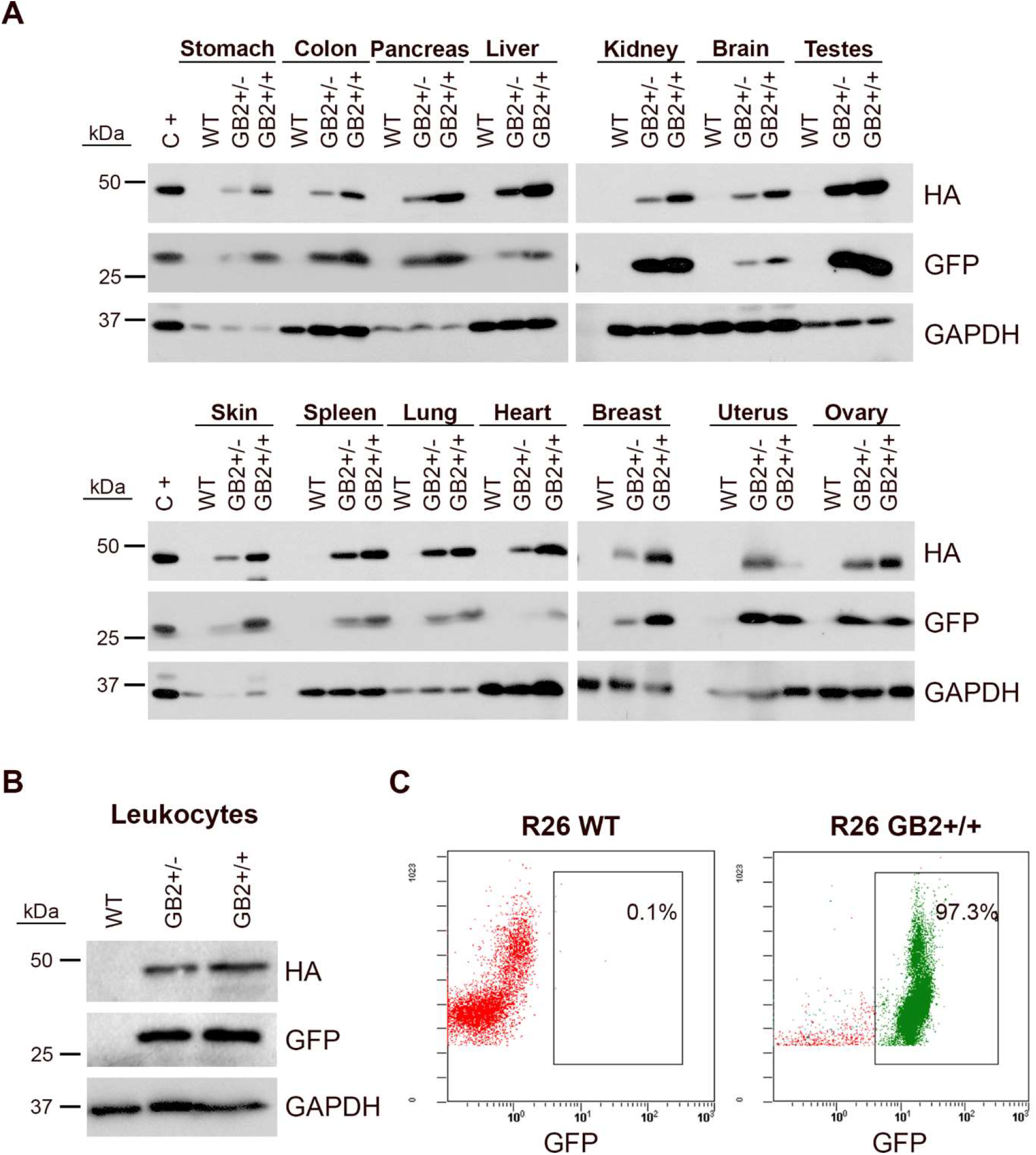
Ubiquitous expression of GSDMB2-HA and GFP in the R26-GB2 mouse model. **A:** Representative western blot analysis in different tissues from GB2 (+/-heterozygous; +/+ homozygous) and WT (control) littermate mice. GAPDH was used as a loading control. C+, MCF7 exogenously expressing GSDMB2-HA and GFP genes were used as a positive control. **B-C:** Expression of GSDMB2-HA and GFP transgenes by WB (B) and GFP by flow cytometry (C) in whole blood leukocytes from R26-GB2 mice. Full-length blots are presented in Supplementary data: Figure S1.

**Fig. 3.**
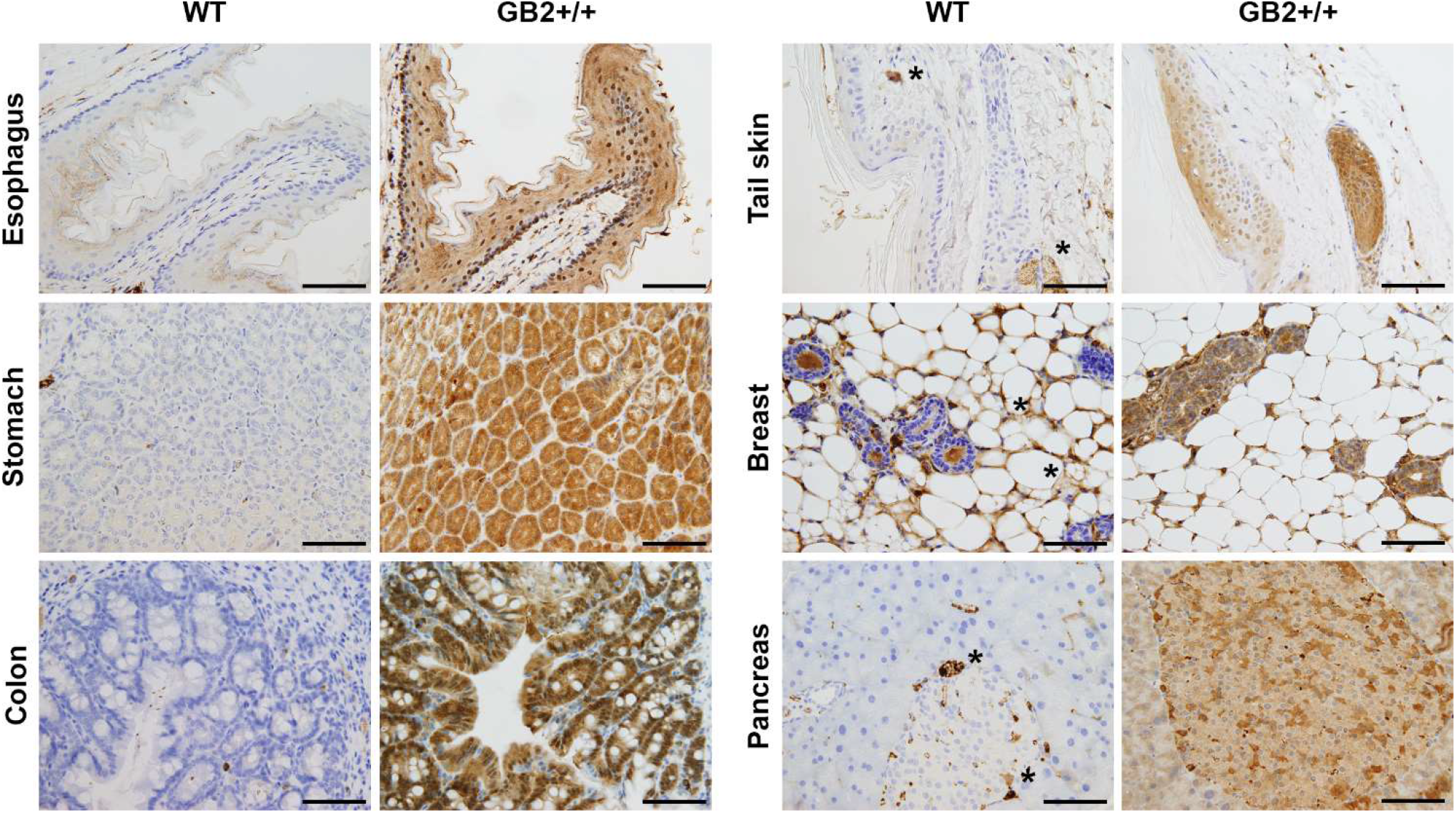
Immunohistochemical expression of GSDMB2-HA in different tissues of the R26-GB2 mouse model. Representative images of tissues from homozygous (GB2+/+) and control (WT) mouse littermates. * Unspecific staining. Scale bar, 100 µm.

To confirm the nuclear-cytoplasmic localization, additional staining with our anti-GSDMB monoclonal antibody [33] was performed in testis. Both HA and GSDMB antibodies showed the same expression pattern (**Supplementary data: Figure S4 A**). Moreover, the nuclear localization in this tissue was confirmed further by immunofluorescence and confocal microscope analysis (**Supplementary data: Figure S4 B**). Overall, the nuclear-cytoplasmic pattern of GSDMB2-HA in mice resembles to that observed in human tissues and cancers [29, 30, 33]. Moreover, the differences found in the intracellular localization pattern among tissues could indicate that GSDMB2 has possibly distinct functions depending on the cell context.

### Effect of GSDMB2 on spontaneous tumorigenesis *in vivo*

Compared to normal tissues, the up-regulation of GSDMB, either at mRNA and protein levels, is commonly seen in multiple tumor types [6, 27-33]. Moreover, GSDMB over-expression promotes multiple pro-tumor functions in breast cancer cells [32-34], in particular the isoform 2 [32]. This data led to the hypothesis that GSDMB could have intrinsic oncogenic properties, but this has not been functionally tested *in vivo*. To investigate if GSDMB2 expression has spontaneous tumorigenic activity in any tissue, or other pathologic consequences, we studied 80 mice for up to 18 months. Mice were monitored weekly for the appearance of tumor masses or other pathological signs and were sacrificed when they showed any of the criteria for early termination specified in Methods or reached 18 months of age. Five mice (all WT) were found dead and necropsy could not be performed, thus post-mortem analyses were done in 75 mice. The overall survival of all the animals (including mice found dead and those sacrificed according to early termination criteria) was similar among GB2+/-, GB2+/+ and WT mice (log-rank Mantel Cox test, p=0.6). At necropsy, tumor formation was investigated in multiple tissues and organs, but macroscopic cancers were only frequently seen in the lungs and stomach (**Table 1**). In fact, the most common neoplasias observed (41 % including all mice) were lung adenocarcinomas, which is consistent with the frequency of these spontaneous tumors in elder mice of the FVB background [47]. However, no significant differences in the frequency of lung tumors (**Table 1**) were observed between WT and GB2 mice (Chi^2^ test p=0.20, considering the three genotypes; and Fisher’s exact test p=0.79 comparing WT versus the combination of GB2+/+ and GB2+/-). Moreover, most of these tumors were well-differentiated lung adenocarcinomas, and no differences in histological grade among the genotypes were observed (**Supplementary data: Table S3**).

**Table 1.** Frequency of spontaneous tumors in GSDMB2-HA knock-in mouse model (R26-GB2) and control (WT) mice.

| GENOTYPE &<br>GENDER | N | LUNG | STOMACH | OTHERS |
| --- | --- | --- | --- | --- |
| WT Male | 13 | 7 (54%) | 3 (23%) |  |
| WT Female | 10 | 2 (20%) | 1 (10%) | 1 Lymphoma |
| **WT TOTAL** | **23** | **9 (39%)** | **4 (17%)** |  |
| GB2+/- Male | 17 | 9 (53%) | 0 |  |
| GB2+/- Female | 16 | 8 (50%) | 0 | 1 Hepatocarcinoma |
| **GB2+/- Total** | **33** | **17 (52%)** | **0** |  |
| GB2+/+ Male | 9 | 2 (22%) | 0 | 1 Lymphoma |
| GB2+/+ Female | 10 | 3 (30%) | 0 | 1 Breast cancer |
| **GB2+/+ Total** | **19** | **5 (26%)** | **0** |  |
| **All mice** | **75** | **31 (41%)** | **4 (5%)** | **4 (5%)** |

GSDMB2 homozygous (GB2+/+), heterozygous (GB2+/-) and WT animals were generated by crossing parental heterozygous mice. Mice were monitored for up to 18 months of age and tissues with macroscopic tumors were analyzed.

Next, to ensure that GSDMB2-HA protein was expressed in these lung tumors, we performed immunohistochemical analyses using an anti-HA antibody. In lungs from GB2+/+ and GB2+/-animals we confirmed the diffuse cytoplasmic staining (and focal nuclear staining in GB2+/+) of GSDMB2-HA in both carcinoma cells and the normal bronchioles (**Fig. 4**). The positive staining with the C-terminal HA tag proves that the full-length GSDMB2 protein is expressed in tumor cells, but it does not have a clear impact on lung cancer development.

**Fig. 4.**
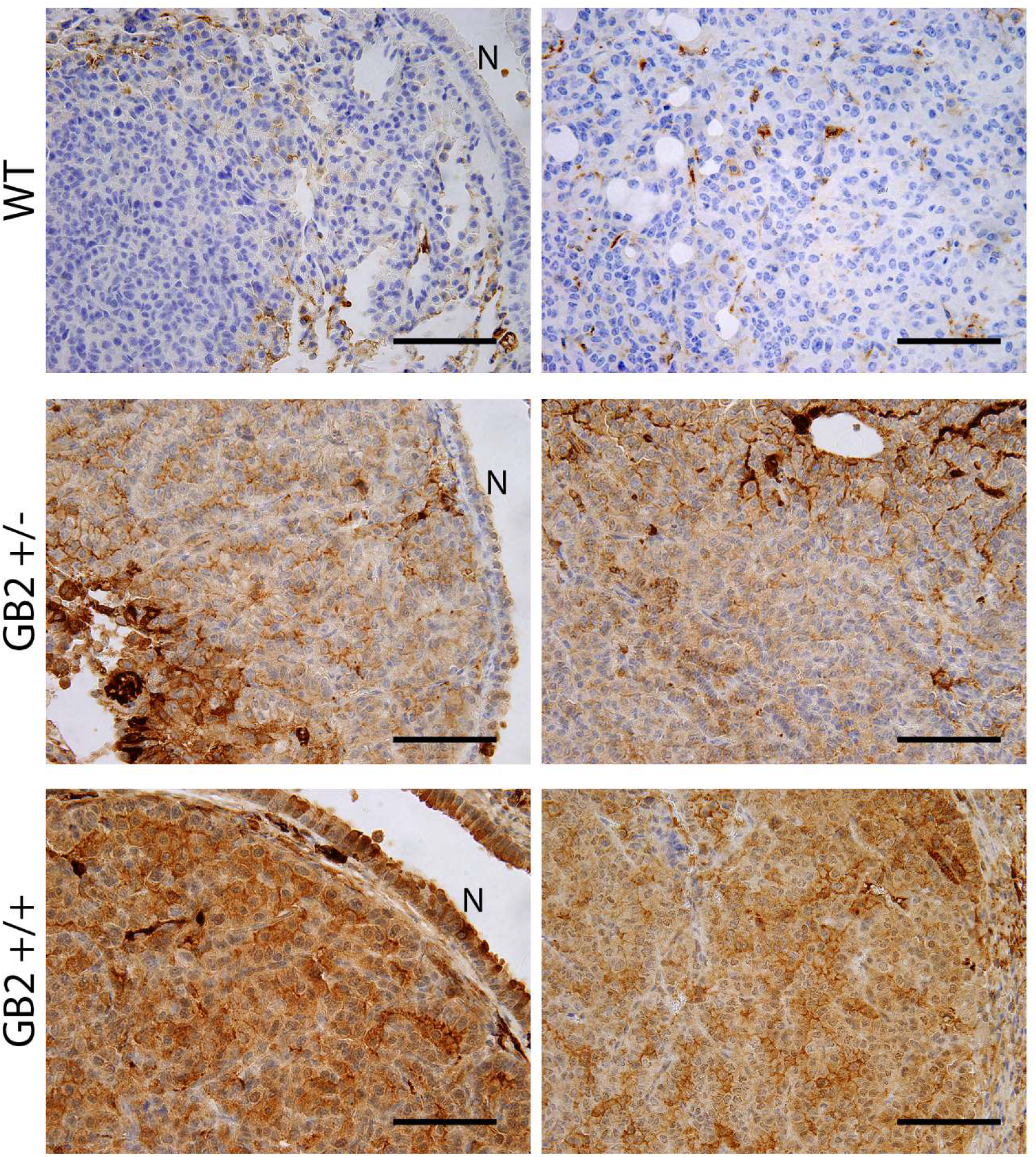
Immunohistochemical expression of GSDMB2-HA in spontaneous lung carcinomas from the R26-GB2 mouse model. Representative images of lung cancers from homozygous (GB2+/+), heterozygous (GB2+/-) and control (WT) mice. Note the stronger expression of GSDMB2-HA in GB2+/+ than GB2+/- cancer cells and the negative staining in the WT condition. N: normal lung bronchiole. Scale bar, 100 µm.

The second most frequent cancer type were gastric tumors. Unexpectedly, we observed that 17% WT mice (3 male and 1 female) developed macroscopic gastric carcinomas, but none of GB2 mice did (**Table 1**) (Chi^2^ test p=0.008, considering the three genotypes; and Fisher’s exact test p=0.007 comparing WT versus the combination of GB2+/+ and GB2+/-). This result is unanticipated, as in humans, GSDMB is generally over- expressed in gastric cancers compared to normal stomach tissues [27-29].

Moreover, while GSDMB over-expression is frequent in human breast cancers [32-34], we only detected one case of spontaneous breast carcinoma in the GB2+/+ mice. Other types of cancer were seldom observed in GB2 or WT animals (**Table 1**), thus, taking all cancers together (irrespective of the tissue of origin) there were no differences in tumor frequency among the mouse genotypes (Chi^2^ test p=0.28, considering the three genotypes; Fisher’s exact test p=0.33 comparing WT versus the combination of GB2+/+ and GB2+/-).

Additionally, since frequent tumors were only seen in lung and stomach, to assess further the effect of GSDMB2 in tumorigenesis, we focused our histological analyses to these organs. Therefore, we evaluated the presence of microscopic tumors or pre-malignant lesions in a series of tissue samples not showing macroscopic evidences of cancer (lung, n=39; stomach, n=30; **Supplementary data: Table S4**). No tumors were detected in these samples, and the frequencies of premalignant adenomatous lung hyperplasia, gastric adenomas/polyps, or chronic gastritis, a potential precursor of stomach cancer [48], were similar in WT and GB2 mice (**Supplementary data: Table S4)**.

As a whole, these data suggest that human GSDMB2 alone does not have a strong overall tumorigenic potential in mice, but it might have instead a potential suppressive effect of gastric carcinogenesis.

### Study of breast tumorigenesis and progression in the R26-GB2/MMTV-PyMT mice

Despite the strong association of GSDMB over-expression and human breast cancer aggressiveness [32-34], the number of mammary carcinomas detected in the R26-GB2 model was scarce. This result suggests that the pro-tumor functions of GSDMB observed in human breast cancers [32-34] may depend on the pre-activation of specific oncogenic stimulus. To test this hypothesis, we evaluated the effect of GSDMB2 expression on breast cancer generation and progression in concert with the PyMT, a strong oncogene [44]. To this end, we generated a double transgenic model, termed R26-GB2/MMTV-PyMT, that expresses GB2 ubiquitously (including the breast) and PyMT specifically in the mammary gland. Breast cancer development was compared between female GB2+/+; PyMT+/- (n=10) and WT; PyMT+/- (n=17) mice. As described before [44], at 15 weeks of age PyMT-driven carcinogenesis provoked the formation of tumors in multiple mammary glands in all animals, thus, tumor incidence was 100% in both GB2+/+ (10/10) and WT (17/17) mice. In GB2+/+ animals we confirmed the strong nucleus-cytoplasmic expression of GSDMB2 in carcinoma cells (**Fig. 5A**). At the histological level, the majority of tumors showed high-grade solid invasive pattern, and no differences were observed between GB2+/+ and WT conditions. Moreover, while cancer latency (time until detection of palpable tumor) was very similar between the conditions (GB2+/+ = 73.8±3.2 days; WT = 73.9 ± 2.3. **Fig 5B**), the number of tumors (per animal) and the tumor weight (either the average per animal or all tumors taken individually) tended to be higher in WT than GB2+/+ mice, though the differences did not reach statistical significance (**Fig. 5**). To test if increased tumor weight in WT mice might reflect an enhanced cancer proliferation, we performed PCNA staining (in the biggest tumor of each mice) but no differences were seen between WT (70.9 ±2.4 percent) compared to GB2+/+ 70.0 ±4.5 percent; **Fig. 5**).

**Fig. 5.**
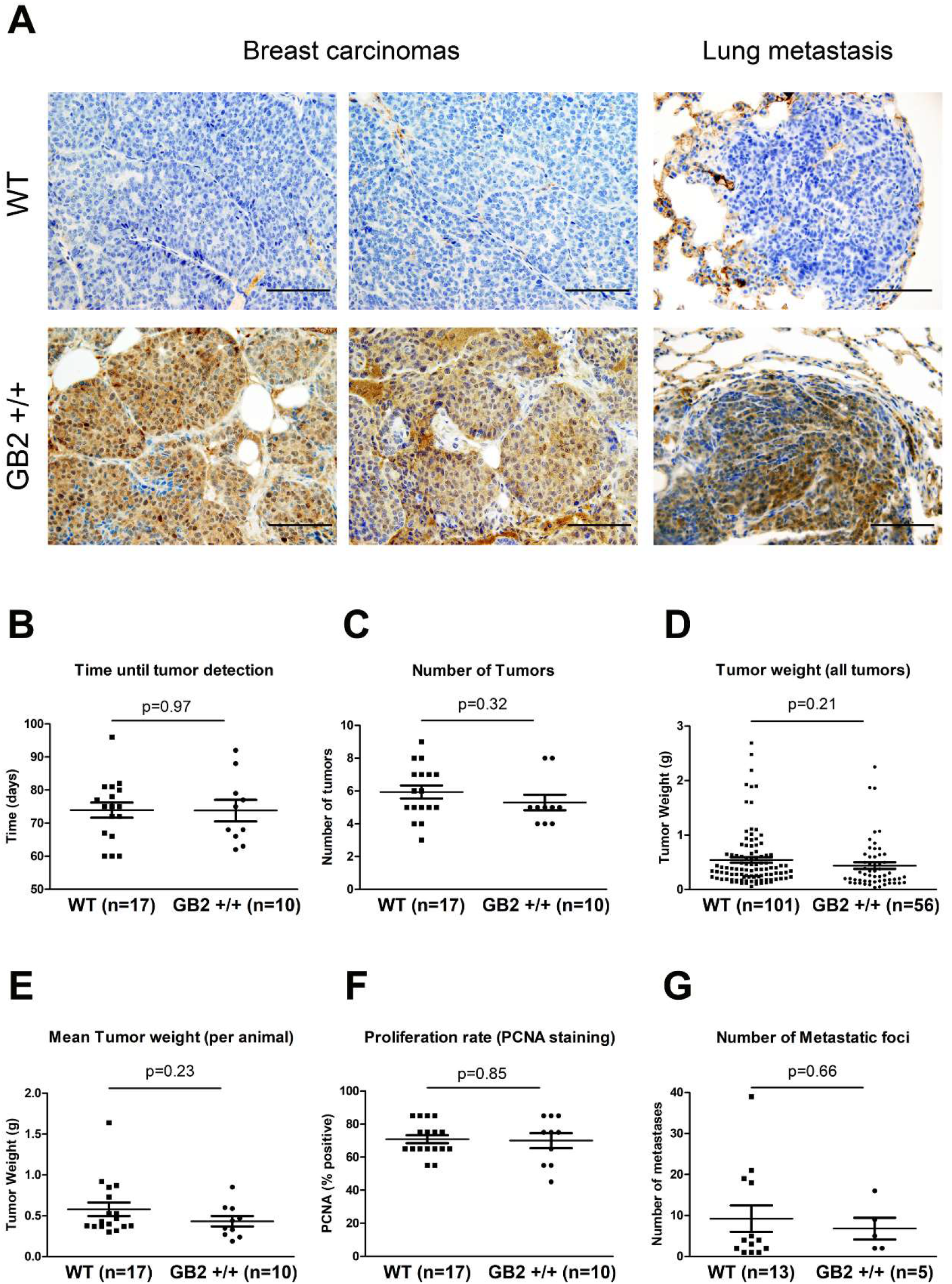
Effect of GSDMB2 on breast cancer generation and progression in the R26-GB2/MMTV-PyMT mouse model. Female double transgenic GB2+/+ (GSDMB2 homozygous); PyMT +/- (heterozygous) mice were compared to WT (GSDMB2 -/-); PyMT +/- animals. **A:** Representative images of the GSDMB2-HA immunohistochemical expression in primary breast carcinomas (left) and lung metastasis (right) from GB2+/+ and WT mice of 15 weeks age. Scale bar, 100 µm. **B:** Comparison of tumor latency (time until detection of palpable mammary tumors). **C:** Average number of breast tumors per animal. **D:** Mean tumor weight of all carcinomas. **E:** Average tumor weight per animal. **F:** Proliferation rate. Percentage of cancer cells positive for PCNA staining. Only the biggest tumor of each mice was analyzed. **G:** Average number of lung metastasis foci (only animals with metastasis). Graphs represent all data and mean values (line). Statistical differences were tested by Student’s t-test.

Next we evaluated the effect of GSDMB2 expression on the metastatic potential. WT mice exhibit more frequently lung metastases (13/17; 76%) than GB2+/+ (5/10; 50%), but differences were not significant (Fisher exact test, p =0.2). The number of metastatic foci (after whole lung sectioning) was very variable within groups, and no clear effect of the mouse genotype was detected (**Fig. 5**). The expression of GSDMB2-HA in lung metastasis was also confirmed by immunohistochemistry (**Fig. 5A**).

Taking together, these results indicate that, contrary to the effect on human breast cancer MCF7 cells [32], GSDMB2 upregulation does not have a clear impact on breast cancer progression in these transgenic mice.

### Analysis of other histopathological alterations in the R26-GB2 model

Finally, apart from cancer, GSDMB has been potentially implicated in the pathobiology of other diseases, including asthma and other inflammatory diseases [6, 37-39, 41, 49-51]. Therefore, in R26-GB2 mice we investigated whether they exhibited any pathological (non-cancer) phenotype at the microscopic level. While lung pathologies (atelectasis and emphysema) tend to be more frequent in GB2 mice than WT, the differences were not statistically significant (**Supplementary data: Table S5**). Moreover, our comprehensive analysis of multiple tissues detected infrequent pathological features in other organs but none of them associated significantly with GSDMB2 expression (**Supplementary data: Table S5**).

## DISCUSSION

Gasdermin proteins play complex and sometimes opposed roles in cancer [6-9, 19-21]. While GSDME and GSDMA are broadly considered as tumor suppressor genes the implication of the other GSDMs in malignancy is less clear [6-9, 19-21]. In particular, GSDMB, which is upregulated in multiple cancers, promotes multiple pro-tumor functions [6, 32-34], and has been considered as a potential oncogene [27]. However, it may also possess “ activatable” cytotoxic anti-tumor effects [34, 35]. To decipher the array of GSDMB *in vivo* functions in cancer it is required the development of genetically engineered mouse models (GEMM). Recently, the group of Dr. Broide reported the first KI model of human GSDMB isoform 3 (the longest isoform) [37]. The hGSDMB^Zp3-Cre^ that ubiquitously expresses GSDMB3 under the control of the strong CAG promoter, shows an asthmatic phenotype associated with increased airway hyper-responsiveness and airway remodeling [37]. Unfortunately, in this model the effect of GSDMB in cancer development was not studied. Here, we analyzed the GSDMB isoform 2, since this transcript increases invasiveness and metastatic potential in breast cancer cells [32]. We generated and characterized phenotypically the first GSDMB2 KI model and tested for the first time whether this protein alone has tumor initiation capacity *in vivo*. After comprehensive analyses of multiple tissues and organs, we proved that GSDMB2 ubiquitous expression in mice neither increases overall tumor development nor affects significantly the aggressiveness of spontaneous generated lung carcinomas. Conversely, only a potential anti-cancer effect was observed in the stomach (discussed later). Given that GSDMB over-expression, in particular the isoform 2, promotes diverse pro-tumor functions in human breast cancers [32-34], we hypothesized that GSDMB functions may require the presence of an oncogenic stimulus. To test this possibility, we evaluated the effect of GSDMB2 expression on breast cancer generation and progression in concert with the PyMT, a strong oncogene in the mammary gland [44]. Surprisingly, in the R26-GB2/MMTV-PyMT model, control mice (lacking GSDMB) tended to generate bigger tumors and more frequently metastatic than GB2+/+ animals, albeit the differences were not statistically significant. In fact, we did not evidence a clear effect of GSDMB2 expression on diverse parameters of cancer formation (number of tumors, latency, tumor weight or cancer histology) or progression (lung metastasis potential). Additionally, in these tumors GSDMB had no impact on cell proliferation, a result consistent with our data in human cancer cells [32, 34].

To ensure that GSDMB2 expression was not lost in cancer cells, in both R26-GB2/MMTV-PyMT and R26-GB2 models we performed immunostaining using an antibody against the C-term HA tag. Results showed that full length GSDMB2 was strongly expressed in both cancer and normal cells (lung or breast). Thus, taken together the data from these models indicate that GSDMB2 upregulation in mice does not have a clear impact on cancer genesis, differentiation and progression. Nonetheless, it is still possible that GSDMB could differentially regulate signaling pathways or affect the function of oncogenes or tumor suppressors in these contexts. However, we did not test this hypothesis since no clear differences were noted in tumor behavior compared with the corresponding control conditions.

Importantly, while our results may not replicate the GSDMB effects observed in human cancer cells, there are a number of potential biological factors that might be required to unveil the full pro-cancer effects of GSDMB in mice: a) in human tumors multiple GSDMB isoforms are co-expressed, that altogether could cooperate in GSDMB cancer activities. b) The levels of GSDMB2 expression in our mouse models might be not high enough to provoke tumorigenic effects. c) GSDMB functions may require the presence of specific stimulus (e.g., pre-activation of precise oncogenes, such as HER2 [33, 34]) or particular cellular contexts. Regarding the biological context, it should be noted that new data showed a potential role for GSDMB as a tumor suppressor, when the immune system is activated [35]. Specifically, Zhou and cols. (2020) reported that GSDMB intrinsic cytotoxic activity in tumor cells could be activated via a non-cell autonomous mechanism mediated by NK and CD4+ T cells [35]. Immunocyte released GZMA cleaves and activates GSDMB thus killing tumor cells *in vitro*. However, *in vivo* models using two aggressive murine cancer xenograft exogenously expressing human full length GSDMB demonstrated that this anti-tumor effect was only possible if additional activation of the immune system was provoked by PD-L1 immune checkpoint inhibitors [35]. This suggests that to trigger an endogenous tumor reduction in mice via the GSDMB cytotoxic mechanism must require additional signals. It is possible that the immune recognition and stimulation of the anti-tumor response may be more difficult to achieve in transgenic animals where the tumor and the surrounding cells carry the same genetic modifications (like our models). In agreement with this idea, Croes et al [52] reported that murine GSDME Knock-out (KO) models did not validate the tumor suppressive effect of GSDME previously observed *in vivo* in xenograft models [15, 21, 53]. While reduced tumoral inflammation was observed in GSDME KO mice, no clear effects on carcinogenesis, tumor differentiation and progression were evidenced using two experimental models of intestinal cancer (chemical induction by azoxymethane or the Apc1638N/+ intestinal cancer mouse strain) [52].

Despite this potential limitation of GSDM GEMM cancer models, our study offers interesting and novel data. First, in the R26-GB2 strain we observed that four WT animals, but no GSDMB2-positive mice developed macroscopic gastric carcinomas. These results were unforeseen, as compared to normal tissue GSDMB is generally over-expressed in human gastric carcinomas [27-29]. In this context, GSDMB upregulation depends on the different usage of the two gene promoters (LTR-derived and cellular promoter) by normal and neoplastic cells [28, 54], but whether GSDMB2 and other isoforms are differentially expressed in cancer and healthy stomach tissue is still unknown. The potential mechanisms by which GSDMB2 might reduce gastric carcinomas could not be explored in our mice as we did not obtain any GSDMB2-positive gastric cancer. Moreover, a potential GSDMB-mediated immune rejection of gastric tumor cells, as described by Zhou et al [35] is unlikely in our model since GSDMB2 is expressed also in immunocytes. Therefore, evaluating further the functional role of GSDMB isoforms in gastric cancer will require future studies using the recently described stomach-specific gene promoters [55] and/or crossing GSDMB models with GEMMs that develop gastric carcinomas [56].

Second, we demonstrated that GSDMB2 shows different nuclear and/or cytoplasmic localization in specific tissues/cell types from healthy organs. Moreover, in both spontaneous lung carcinomas (R26-GB2 strain) and PyMT-driven breast cancers (R26-GB2/MMTV-PyMT) GSDMB2 staining was mostly cytoplasmic but nuclear localization was noted in some tumor areas. All these data suggest that this protein may have distinct biological effects depending on the cellular context or microenvironment. Similarly, in human normal and cancer tissues GSDMB has been observed in the cytoplasm and the cell nucleus [27, 29, 30, 33]. GSDMB possesses a sequence similar to a nuclear localization signal (residues 242-261), encoded by exon 8, that is present in all GSDMB isoforms, and mutation/deletion of this sequence excludes GSDMB from the nucleus [30, 37]. Despite to date no mechanism of the GSDMB nucleus-cytoplasm shuttling has been reported and the biological function of nuclear GSDMB localization is unclear, some evidences indicate that GSDMB may indirectly regulate transcription of specific genes. Consistent with this, in human bronchial epithelial cells, nuclear accumulation of GSDMB isoform 1 is required for the transcriptional induction of TGF-β1 and 5-lipoxygenase [37]. Interestingly, in the mouse hGSDMB^Zp3-Cre^ model upregulation of the same genes also occurred and led to airway remodeling. These results suggest that human GSDMB is able to produce biological effects by modulating specific transcripts even in the mouse genome. In this sense, our novel mouse models will be useful in future studies to assess whether GSDMB regulates specific genes in particular tissues/cell types from both healthy and cancer conditions.

Third, apart from cancer, the GSDMs have been functionally linked to a variety of diseases ranging from septic shock (GSDMD) to deafness syndromes (GSDME & PJVK), among others [3-5]. Specifically, GSDMB has been associated to multiple inflammatory pathologies, such as asthma, type-I diabetes, inflammatory bowel diseases, biliary cirrhosis, rheumatoid arthritis, and idiopathic inflammatory myopathies [6, 39, 41, 49-51]. However, the functional implication of GSDMB has only been demonstrated for asthma, both in human cells [38] and transgenic mice [37]. Interestingly, GSDMB isoforms may play different roles in this disease [37-39]. While we did not evaluate the presence of asthmatic phenotype in our R26-GB2 model, we observed that other lung pathologies (atelectasis and emphysema) tend to be more frequent (albeit not significant) in GB2 mice. Future studies comparing GSDMB3 and GSDMB2 mouse models in lung disease would be of interest to confirm these findings. Apart from lung, in our comprehensive histological examination of multiple organs from R26-GB2 mice we did not observe any frequent and consistent pathological alteration in the tissues studied. However, as discussed before, it should be noted that GSDM functions, in particular their pro-cell death function, are activated under specific circumstances. In fact, some GEMMs of GSDMs reported to date exhibit pathological phenotypes only when they carry gain-of-function mutations or in response to specific stimuli. For instance, *Gsdma1* KO mice shows no phenotype, but *Gsdma1* KI over-expressing and KI mutant (A339T) mice display epidermal hyperplasia [57]. *Gsdmd* KO mice do not show abnormalities in the digestive system [58] but those animals are completely resistant to septic shock (pyroptosis-mediated) induced by LPS injection [13].

All these data suggest that to fully decipher the role of GSDMs in cancer and other pathologies using GEMMs it is first required to identify the precise stimulus and molecular mechanisms governing their biological functions in humans.

## CONCLUSIONS

The phenotypic characterization of our novel knock-in models indicates that nucleus and/or cytoplasmic GSDMB2 expression alone does not have an overall tumorigenic effect in mice. Moreover, in spontaneous lung carcinomas or PyMT-driven breast cancers, GSDMB2 upregulation does not have a clear impact on cancer behavior, differentiation and progression. Nonetheless, we observed a potential reduction of gastric carcinogenesis in GSDMB2-positive mice that requires further study. It should be noted that unveiling the array of GSDMB *in vivo* functions in cancer may require the presence of specific stimulus or particular cellular contexts. Moreover, it will be of great interest to generate and compare models expressing the distinct GSDMB isoforms for assessing the importance of these protein variants in the development and progression of cancer and other diseases. In all these aspects, our new models will serve as the basis for the future development of more precise tissue-specific and context-dependent cancer models.

## LIST OF ABBREVIATIONS

aas: Aminoacids
GB2: Gasdermin-B isoform 2
GFP: Green Fluorescent Protein
GSDM: Gasdermin
GEMM: Genetically engineered mouse models
KO: Knock-out
KI: Knock-in
PyMT: Polyoma Middle T antigen
R26: Rosa26 locus
WB: Western Blot
WT: Wildtype

## DECLARATIONS

### Ethics approval and consent to participate

All mouse studies were performed in agreement with the procedures and protocols that have been approved by the internal committees of ethical and animal welfare of the Institutions (Autonomous University of Madrid and Institute of Biomedical Sciences Alberto Sols-CSIC) and the local authorities (Comunidad de Madrid, PROEX424/15). The procedures comply with the European Union (Directive 2010/63/UE) and Spanish Government guidelines (Real Decreto 53/20133). All animals were housed in the IIBm animal facility within the same room under standard conditions, a maximum of 4 animals per cage. No animals were caged individually. The study reporting adheres to the ARRIVE guidelines, and a completed ARRIVE checklist is provided as **Supplementary data: File 6**.

## Consent for publication

Not applicable. This work does not involve human studies.

## Availability of data and materials

All data generated or analyzed during this study are included in this article and its supplementary information files.

## Authors’ contributions

DS conceived and designed the study, contributed to the generation and characterization of the mouse models, performed the experiments, generated and analyzed the data and wrote the manuscript. ARS and AT performed the histopathological analyses of the tissue samples. MPL performed WB experiments, contributed to generation of *in vivo* data and discussed the results. EDM performed histological and immunohistochemical staining. LMS collected tissue samples, performed WBs and the mouse genotyping. PGS generated the targeting vector. JP contributed to tissue sample processing and histological analyses. GMB conceived, designed and directed the study, generated and analyzed the data, contributed to histological and immunohistochemical studies and wrote the manuscript. All the authors read and approved the final manuscript.

## Acknowledgements

The authors would like to thank all current and past members of the Moreno-Bueno’s lab and Amparo Cano’s group, in particular Dr. Alberto Martin for their invaluable help in the development of the mouse models. We are grateful to the scientific units involved in this project: IIBm Animal facility, CNIO Transgenic Mice Unit and MD Anderson Cancer Center Pathology Lab.

