## Supplementary data for "IN VIVO ANALYSIS OF THE ROLE OF GASDERMIN-B (GSDMB) IN CANCER USING NOVEL KNOCK-IN MOUSE MODELS"

Sarrio D et al.

**Table S1: Primers and PCR conditions for genotyping**

| PRIMERS | bp | PCR conditions |
|---|---|---|
| **Transgene insertion into ROSA26 (3'arm)**<br>3'DV1-F1: 5'-GAACGGCATCAAGGTGAAC-3'<br>3'DV1-R1: 5'-ATCTCGAAGACCTGTTGCTG-3' | 4800 | 94ºC 1min; (95ºC 30s, 64ºC 30s, 68ºC 5min) X40; 72ºC 7min. LATaq Polymerase (Takara) |
| **Transgene insertion into ROSA26 (5'arm)**<br>NeoF3: 5'-CCAGTCATAGCCGAATAGCC-3'<br>5'DV1-F2: 5'-TAGGTAGGGGATCGGGACTC-3' | 1800 | 94ºC 1min; (94ºC 30s, 68ºC 150s) X5; (95ºC 30s, 64ºC 30s, 68ºC 135s) X 35; 72ºC 7min. LATaq Polymerase (Takara) |
| **CRE**<br>CreF: 5'-GCCTGCATTACCGGTCGATGC -3'<br>CreR: 5'-CAGGGTGTTATAAGCAATCCCC -3' | 430 | 94ºC 5min; (94ºC 30s, 60ºC 30s, 72ºC 30s) X35; 72ºC 5min. NZYTaq II Polymerase (NZYTech) |
| **Excision of NEO cassette & GSDMB2 activation**<br>GS-CreF: 5'-TATCAGTAAGGGAGCTGCAGTG-3'<br>GS-CreR3: 5'-TGAGGCCTGTTGTGTAGTGC-3' | 700 | 94ºC 5min; (94ºC 30s, 60ºC 1 min, 72ºC 1min) X35; 72ºC 2min. NZYTaq II Polymerase (NZYTech) |
| **ROSA26 WT allele**<br>ROSA26-F: 5'-TATCAGTAAGGGAGCTGCA -3'<br>ROSA26-R: 5'-ACCCCAGATGACTACCTATCC -3' | 300 | 94ºC 5min; (94ºC 30s, 62ºC 30s, 72ºC 30s) X35; 72ºC 5min. NZYTaq II Polymerase (NZYTech) |
| **GSDMB2-HA cDNA**<br>G2HA-F: 5'-TGGGTTCGGAGGATTCCAGA-3<br>G2HA-R: 5'-AGCATAATCAGGAACATCATACGG-3' | 548 | 94ºC 5min; (94ºC 30s, 64ºC 30s, 72ºC 40s) X35; 72ºC 5min. NZYTaq II Polymerase (NZYTech) |
| **GFP cDNA**<br>GFP-F: 5'-AAGGACGACGGCAACTACAAG-3'<br>GFP-R: 5'-AGGTAGTGGTTGTCGGGCAG-3' | 300 | 94ºC 5min; (94ºC 30s, 68ºC 30s, 72ºC 30s) X35; 72ºC 5min. NZYTaq II Polymerase (NZYTech) |
| **PYMT**<br>POL F: 5'-ATCGGGCTCAGCAACACAAG-3'<br>POL R: 5'-AACGGCGGAGCGAGGAACTG-3' | 280 | 94ºC 5min; (94ºC 30s, 61ºC 30s 72ºC 30s) X30 72ºC 5min. NZYTaq II Polymerase (NZYTech) |

**Table S2: Immunohistochemical expression and intracellular localization of GSDMB2-HA in selected tissues from the R26-GB2 mouse model**

| Tissue | GSDMB2-HA staining (intensity and localization) |
|---|---|
| **Skin (tail)** | Strong nuclear and cytoplasmic in epidermis, hair follicles and sebaceous glands. |
| **Esophagus** | Strong nuclear staining in the squamous epithelium. |
| **Stomach** | Moderate cytoplasmic staining in the epithelium. |
| **Intestine** | Strong cytoplasmic staining and focal nuclear localization in the epithelium. |
| **Liver** | Weak cytoplasmic staining in hepatocytes. |
| **Pancreas** | Moderate cytoplasmic staining in pancreatic cells. |
| **Kidney** | Weak cytoplasmic in glomeruli and tubules. Strong cytoplasmic in renal papilla. |
| **Lung** | Weak cytoplasmic and focal nuclear staining in bronchus/bronchioles. |
| **Heart** | Weak cytoplasmic staining in muscle cells. |
| **Brain** | Weak cytoplasmic and focal nuclear overall. Strong cytoplasmic in ependymal cells of choroid plexus. |
| **Breast** | Strong cytoplasmic in mammary gland epithelia. |
| **Salivary gland** | Weak cytoplasmic in glandular cells. |
| **Spleen** | Weak diffuse cytoplasmic staining in lymphoid cells. |
| **Uterus** | Strong cytoplasmic and focal nuclear expression in cervix epithelium. |
| **Testicles** | Very strong cytoplasmic and nuclear staining in seminiferous tubules. Weak cytoplasmic in epididymis. |

**Table S3. Histological characteristics of the spontaneous tumors originated in GSDMB2-HA knock-in model (R26-GB2) and control (WT) mice.**

| TUMOR HISTOLOGY | WT | GB2+/- | GB2+/+ | P value[1] | P value[2] |
|---|---|---|---|---|---|
| **Lung Adenocarcinoma (n)** | **9** | **17** | **5** | 0.69 | 0.59 |
| Well differentiated | 5 (56%) | 11 (65%) | 2 (40%) | | |
| Moderately differentiated | 2 (22%) | 5 (29%) | 2 (40%) | | |
| Poorly differentiated | 2 (22%) | 1 (6%) | 1 (20%) | | |
| **Gastric carcinoma (n)** | **4** | **0** | **0** | ND | ND |
| Low grade | 3 (75%) | 0 | 0 | | |
| High grade | 1 (25%) | 0 | 0 | | |

GSDMB2 Heterozygous (GB2+/-), homozygous (GB2+/+) and control (WT) animals were generated by crossing parental heterozygous mice. [1] p value of $Chi^2$ test comparing the three genotypes separately; [2] p value of Fisher's exact test comparing WT vs GB2 (+/- and +/+ combined). ND, not done

**Table S4. Pre-malignant microscopic lesions in lungs and stomach from GSDMB2-HA knock-in model (R26-GB2) and control (WT) mice.**

| Lesion | WT | GB2+/- | GB2+/+ | P Value[1] | P Value[2] |
|---|---|---|---|---|---|
| Gastric adenomas and polyps | 1/7 (14%) | 0/13 (0%) | 1/10 (10%) | 0.4 | 0.41 |
| Chronic gastritis | 3/7 (43%) | 4/13 (30%) | 3/10 (30%) | 0.8 | 0.36 |
| Lung adenomatous hyperplasia | 1/11 (9%) | 0/13 (0%) | 2/14 (14%) | 0.4 | 0.99 |

GSDMB2 Heterozygous (GB2+/-), homozygous (GB2+/+) and control (WT) animals were generated by crossing parental heterozygous mice. [1] p value of $Chi^2$ test comparing the three genotypes separately; [2] p value of Fisher's exact test comparing WT vs GB2 (+/- and +/+ combined).

**Table S5. Frequency of other non-cancer microscopic lesions from GSDMB2-HA knock-in model (R26-GB2) and control (WT) mice.**

| Non-cancer lesion | WT | GB2+/- | GB2+/+ | P Value[1] | P Value[2] |
|---|---|---|---|---|---|
| Lung Emphysema | 1/11 (9%) | 4/13 (31%) | 1/14 (7%) | 0.18 | 0.65 |
| Lung Atelectasis | 0/11 (0%) | 2/13 (15%) | 4/14 (29%) | 0.15 | 0.15 |
| Liver Steatosis or necrosis | 3/5 (60%) | 7/8 (87.5%) | 3/6 (50%) | 0.29 | 0.99 |
| Uterine/ovarian benign cysts | 0/6 (0%) | 4/9 (45%) | 1/5 (20%) | 0.14 | 0.26 |
| Other analyzed tissues with <2 cases of pathology * | 27 | 28 | 27 | | |

GSDMB2 Heterozygous (GB2+/-), homozygous (GB2+/+) and control (WT) animals were generated by crossing parental heterozygous mice.[1] p value of $Chi^2$ test comparing the three genotypes separately; [2] p value of Fisher's exact test comparing WT vs GB2 (+/- and +/+ combined). *: brain, salivary gland, heart, bladder, kidney, intestine, pancreas, spleen and male reproductive organs.

**SUPPLEMENTARY FIG 1:** UNCROPPED GELS FOR FIGURE 1

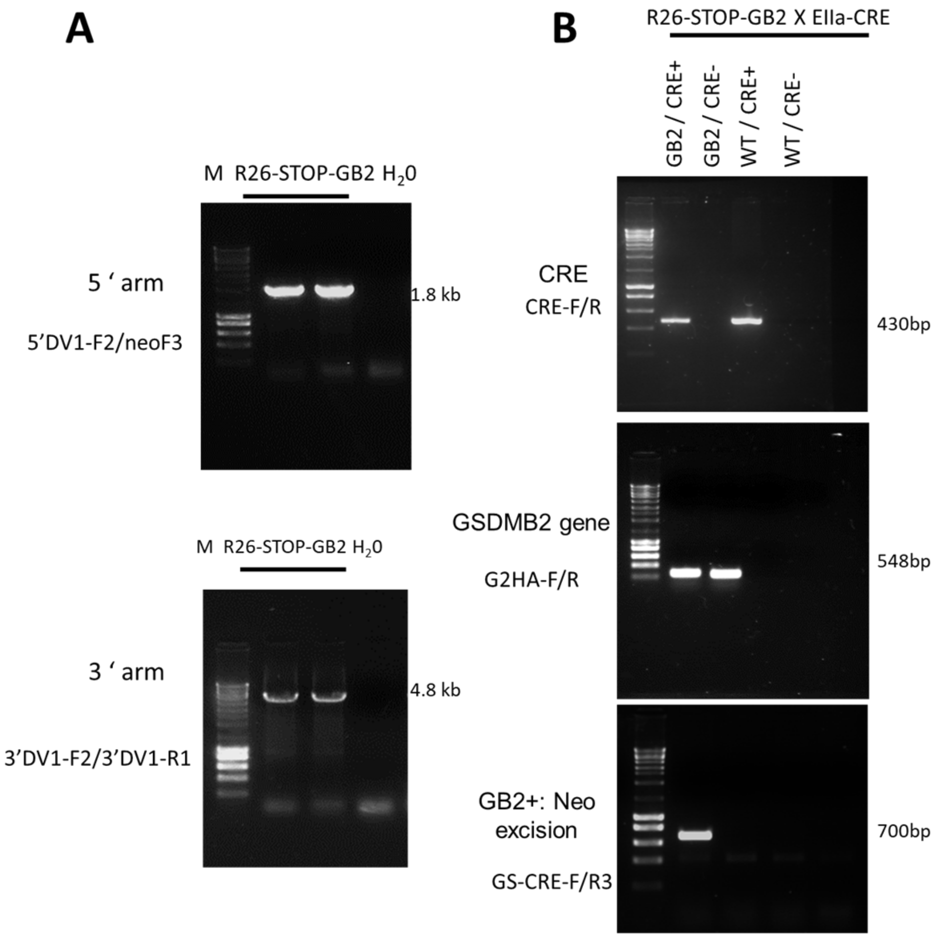

**SUPPLEMENTARY FIG 1:** UNCROPPED BLOTS FOR FIGURE 2 (PANEL A, TOP)

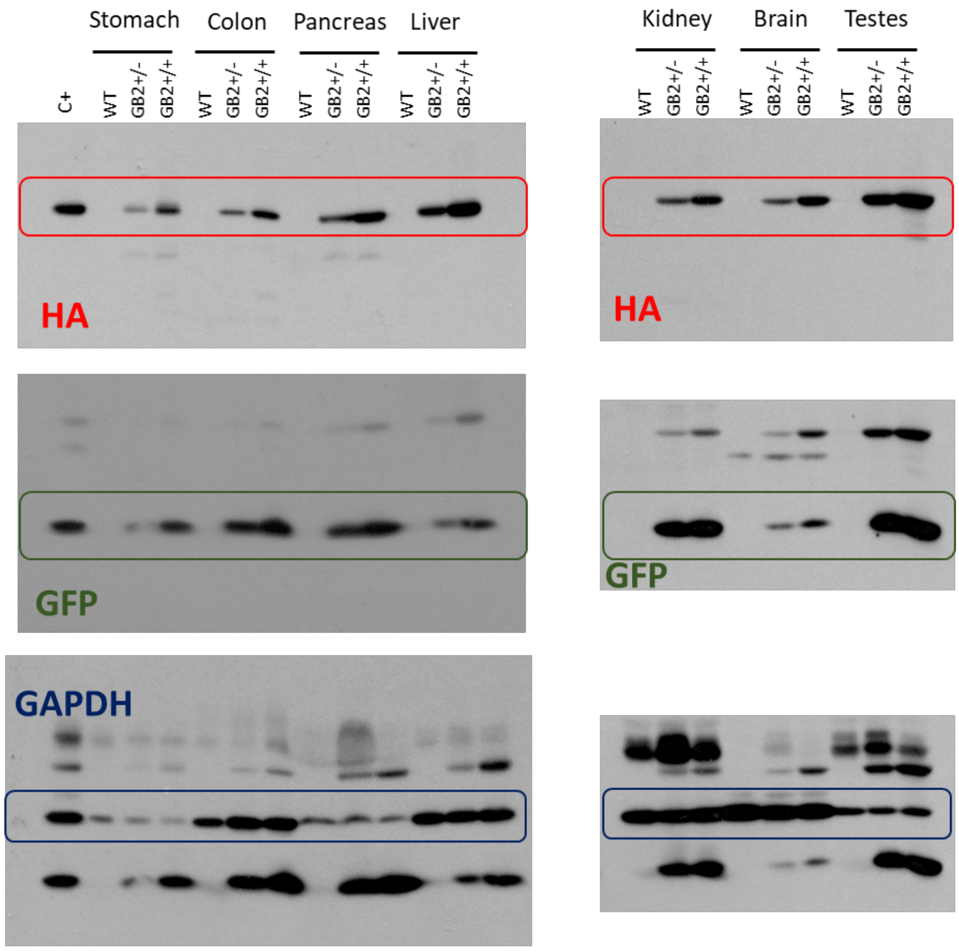


Cropped areas are indicated with color boxes

**SUPPLEMENTARY FIG 1:** UNCROPPED BLOTS FOR FIGURE 2 (PANEL A, BOTTOM)

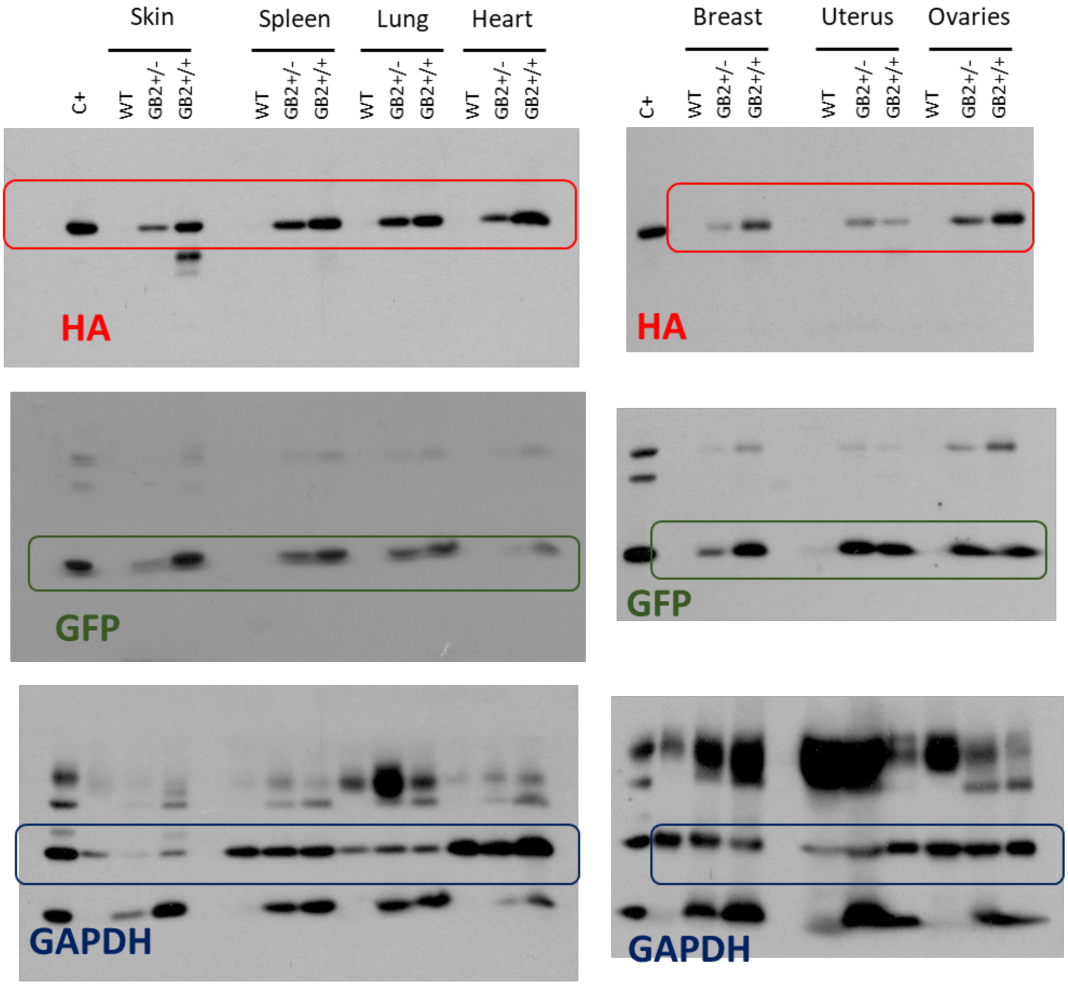


UNCROPPED BLOTS FOR FIGURE 2 (PANEL B)

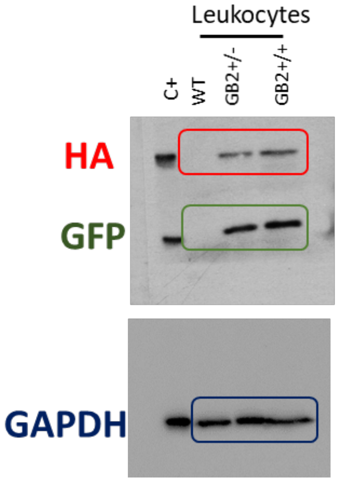


Cropped areas are indicated with color boxes

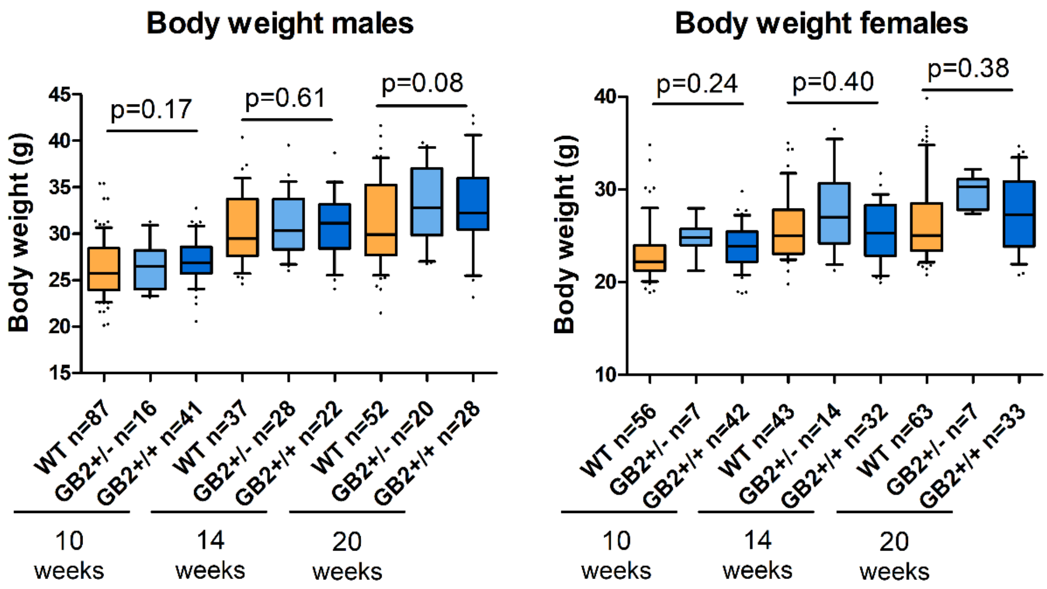


**Supplementary Figure S2. Total body weight of R26-GB2 mice compared to WT controls.** The age (in weeks) and the number of mice per group are indicated. Box plots represent median values and quartiles, and whiskers the 10-90 percentile. Heterozygous (GB2+/-), homozygous (GB2+/+) and control (WT) were generated by crossing parental heterozygous mice. P value of t-test comparing WT and GB2+/+.

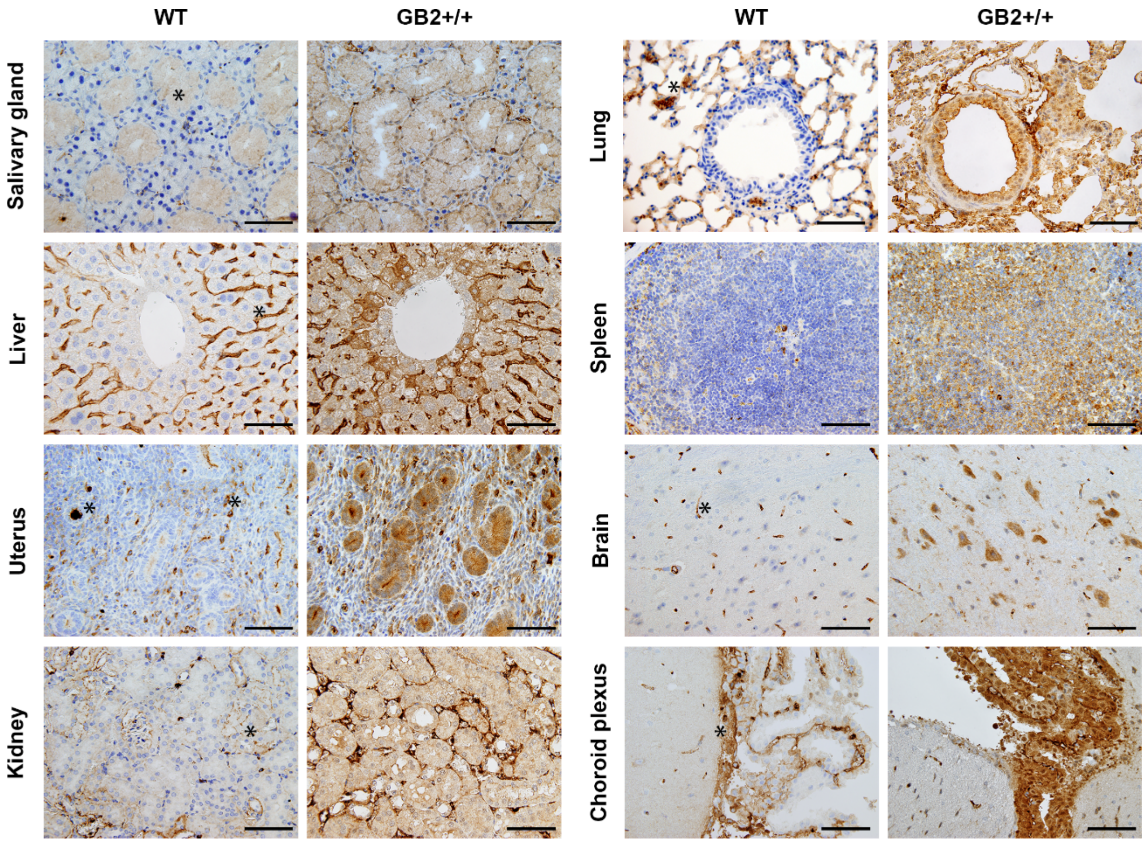


**Supplementary Figure S3. Immunohistochemical expression of GSDMB2-HA in different tissues from the R26-GB2 mouse model.** Representative images of tissues from homozygous (GB2+/+) and control (WT) mouse littermates. * Unspecific staining. Scale bar, 100 µm.

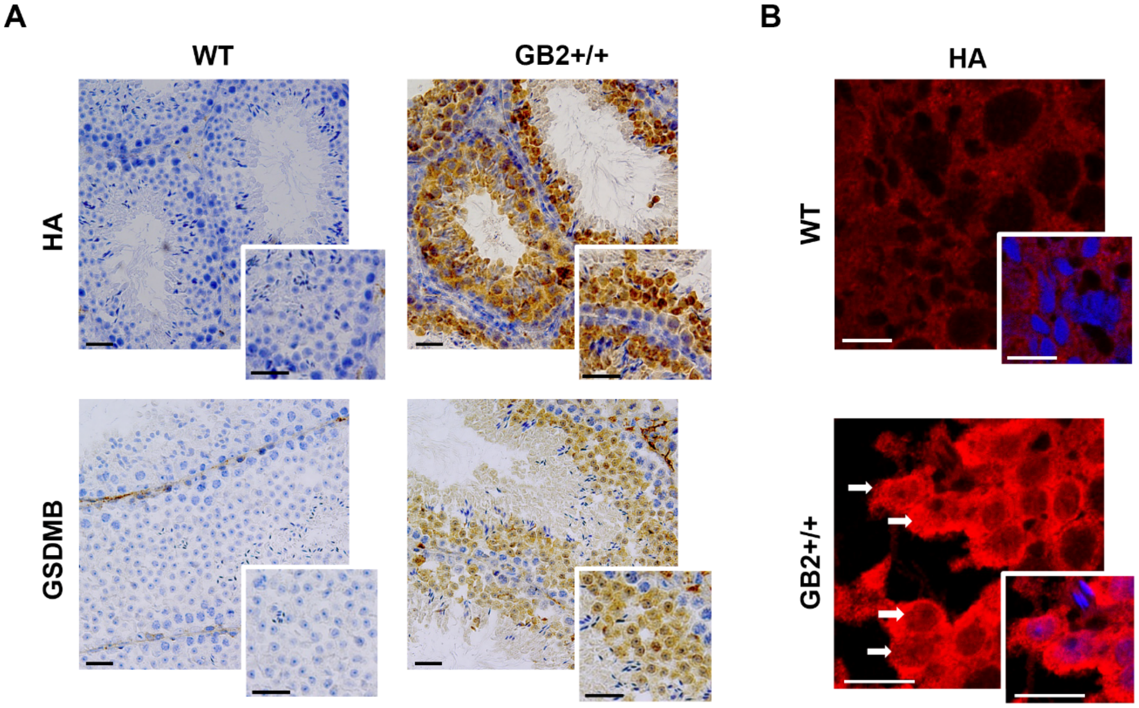


**Supplementary Figure S4. Nuclear and cytoplasmic localization of GSDMB2-HA in the testes of R26-GB2 mice**. A: Immunohistochemical expression using rat anti-HA antibody (top) and mouse anti-GSDMB antibody [33] in testes from WT and homozygous (GB2 +/+) mice. Inset: zoomed images. Scale bar 50 µm. B: Immunofluorescence staining and confocal imaging of GSDMB2-HA in the samples depicted in (A). GSDMB2-HA (red) localizes focally in the cell nucleus (arrows). In the inset images, nuclei are stained with DAPI (blue). Scale bar 25 µm

**Additional File 6: completed ARRIVE checklist (pdf).**

### The ARRIVE guidelines 2.0: author checklist

#### The ARRIVE Essential 10

These items are the basic minimum to include in a manuscript. Without this information, readers and reviewers cannot assess the reliability of the findings.

| Item | | Recommendation | Section/line number, or reason for not reporting |
|---|---|---|---|
| **Study design** | 1 | For each experiment, provide brief details of study design including:<br>a. The groups being compared, including control groups. If no control group has been used, the rationale should be stated.<br>b. The experimental unit (e.g. a single animal, litter, or cage of animals). | |
| **Sample size** | 2 | a. Specify the exact number of experimental units allocated to each group, and the total number in each experiment. Also indicate the total number of animals used.<br>b. Explain how the sample size was decided. Provide details of any *a priori* sample size calculation, if done. | |
| **Inclusion and exclusion criteria** | 3 | a. Describe any criteria used for including and excluding animals (or experimental units) during the experiment, and data points during the analysis. Specify if these criteria were established *a priori*. If no criteria were set, state this explicitly.<br>b. For each experimental group, report any animals, experimental units or data points not included in the analysis and explain why. If there were no exclusions, state so.<br>c. For each analysis, report the exact value of *n* in each experimental group. | |
| **Randomisation** | 4 | a. State whether randomisation was used to allocate experimental units to control and treatment groups. If done, provide the method used to generate the randomisation sequence.<br>b. Describe the strategy used to minimise potential confounders such as the order of treatments and measurements, or animal/cage location. If confounders were not controlled, state this explicitly. | |
| **Blinding** | 5 | Describe who was aware of the group allocation at the different stages of the experiment (during the allocation, the conduct of the experiment, the outcome assessment, and the data analysis). | |
| **Outcome measures** | 6 | a. Clearly define all outcome measures assessed (e.g. cell death, molecular markers, or behavioural changes).<br>b. For hypothesis-testing studies, specify the primary outcome measure, i.e. the outcome measure that was used to determine the sample size. | |
| **Statistical methods** | 7 | a. Provide details of the statistical methods used for each analysis, including software used.<br>b. Describe any methods used to assess whether the data met the assumptions of the statistical approach, and what was done if the assumptions were not met. | |
| **Experimental animals** | 8 | a. Provide species-appropriate details of the animals used, including species, strain and substrain, sex, age or developmental stage, and, if relevant, weight.<br>b. Provide further relevant information on the provenance of animals, health/immune status, genetic modification status, genotype, and any previous procedures. | |
| **Experimental procedures** | 9 | For each experimental group, including controls, describe the procedures in enough detail to allow others to replicate them, including:<br>a. What was done, how it was done and what was used.<br>b. When and how often.<br>c. Where (including detail of any acclimatisation periods).<br>d. Why (provide rationale for procedures). | |
| **Results** | 10 | For each experiment conducted, including independent replications, report:<br>a. Summary/descriptive statistics for each experimental group, with a measure of variability where applicable (e.g. mean and SD, or median and range).<br>b. If applicable, the effect size with a confidence interval. | |

### The Recommended Set

These items complement the Essential 10 and add important context to the study. Reporting the items in both sets represents best practice.

| Item | | Recommendation | Section/line number, or reason for not reporting |
|---|---|---|---|
| **Abstract** | 11 | Provide an accurate summary of the research objectives, animal species, strain and sex, key methods, principal findings, and study conclusions. | |
| **Background** | 12 | a. Include sufficient scientific background to understand the rationale and context for the study, and explain the experimental approach.<br>b. Explain how the animal species and model used address the scientific objectives and, where appropriate, the relevance to human biology. | |
| **Objectives** | 13 | Clearly describe the research question, research objectives and, where appropriate, specific hypotheses being tested. | |
| **Ethical statement** | 14 | Provide the name of the ethical review committee or equivalent that has approved the use of animals in this study, and any relevant licence or protocol numbers (if applicable). If ethical approval was not sought or granted, provide a justification. | |
| **Housing and husbandry** | 15 | Provide details of housing and husbandry conditions, including any environmental enrichment. | |
| **Animal care and monitoring** | 16 | a. Describe any interventions or steps taken in the experimental protocols to reduce pain, suffering and distress.<br>b. Report any expected or unexpected adverse events.<br>c. Describe the humane endpoints established for the study, the signs that were monitored and the frequency of monitoring. If the study did not have humane endpoints, state this. | |
| **Interpretation/ scientific implications** | 17 | a. Interpret the results, taking into account the study objectives and hypotheses, current theory and other relevant studies in the literature.<br>b. Comment on the study limitations including potential sources of bias, limitations of the animal model, and imprecision associated with the results. | |
| **Generalisability/ translation** | 18 | Comment on whether, and how, the findings of this study are likely to generalise to other species or experimental conditions, including any relevance to human biology (where appropriate). | |
| **Protocol registration** | 19 | Provide a statement indicating whether a protocol (including the research question, key design features, and analysis plan) was prepared before the study, and if and where this protocol was registered. | |
| **Data access** | 20 | Provide a statement describing if and where study data are available. | |
| **Declaration of interests** | 21 | a. Declare any potential conflicts of interest, including financial and non-financial. If none exist, this should be stated.<br>b. List all funding sources (including grant identifier) and the role of the funder(s) in the design, analysis and reporting of the study. | |
